# Molecular characterization of MRI/CYREN reveals the Ku binding mode and the role of multimerization in stimulating the activity of NHEJ in DNA repair

**DOI:** 10.64898/2026.07.22.740027

**Authors:** Duc-Duy Vu, Kaela Makins, Sonja Knödlstorfer, Philippe Pelupessy, Ludovic Carlier, Guillaume Bouvignies, Jeremy M. Stark, Mauro Modesti, Fabien Ferrage

## Abstract

Mammalian cells primarily repair DNA double-strand breaks through non-homologous end joining (NHEJ), a pathway that requires the Ku heterodimer, DNA-PKcs, XRCC4 complexed with DNA Ligase 4, and XLF as core components. In addition, several auxiliary proteins are involved in the regulation of NHEJ, whose importance has been underscored recently. Among them, MRI, also known as CYREN, is one of the most important auxiliary proteins. Despite its importance, the structural properties of MRI remain poorly characterized. In this study, we used solution NMR spectroscopy combined with cellular experiments to investigate two isoforms of human MRI (MRI1 and MRI2) at the residue level. Our findings reveal that both isoforms are predominantly disordered, and that the APLF-like Ku-binding motif (A-KBM) of MRI undergoes folding upon binding to the von Willebrand A domain of Ku80 (Ku80_vWA_). Moreover, an evolutionarily dominant leucine-to-methionine substitution in A-KBM significantly increases binding affinity for Ku80_vWA_ by over 30 times without impacting cellular NHEJ efficiency. Importantly, we identified here a domain that drives MRI multimerization that is required for efficient NHEJ in cellular assays. This work further deciphers the increasingly recognized functional roles of disordered protein regions of the NHEJ machinery.

## INTRODUCTION

DNA double-strand breaks (DSB) arise both from exogenous sources, such as ionizing radiation, and from endogenous sources, including replication errors or reactive oxygen species generated during oxidative metabolism (1). Beyond their genotoxic threat, DSBs serve essential physiological functions as programmed intermediates in genome rearrangements during lymphocyte development and adaptive immunity (1, 2). Failure to accurately repair these lesions degrades genomic integrity, predisposing cells to chromosomal rearrangements that can drive tumorigenesis or trigger cell death (1). Therefore, mammalian cells rely on several dedicated DSB repair pathways, of which non-homologous end joining (NHEJ) is the predominant mechanism in human cells (2–4).

Recent structural and single-molecule studies of key stages of the NHEJ pathway have revealed the orchestrated actions of its core components: the Ku70/80 heterodimer (Ku), DNA-dependent protein kinase catalytic subunit (DNA-PKcs), XLF (also called NHEJ1 or Cernunnos), and the XRCC4–DNA ligase 4 complex (X4L4) (3, 5–9). In the consensus model, Ku and DNA-PKcs initially bind and protect the DNA ends, and initiate long-range synaptic complex formation with XLF and X4L4. Subsequently a transition is believed to occur, giving rise to a short-range complex presumably after DNA-PKcs eviction that juxtaposes and ligates the two DNA ends (3, 5–9). We recently demonstrated that the disordered regions of XLF and XRCC4 are also critical in regulating NHEJ activity (10).

In addition to NHEJ core components, several auxiliary factors also participate in regulating NHEJ activity, including Aprataxin and PNK-like factor (APLF), Paralog of XRCC4 and XLF (PAXX), and Modulator of retrovirus infection homolog (MRI, also known as CYREN, Cell cycle regulator of NHEJ). APLF stabilizes the long-range complex (11), whereas PAXX, which is structurally and functionally redundant with XLF, reinforces the NHEJ synapsis through its conserved Ku-binding motif (KBM) that interacts with Ku70 (12–16). The function of MRI is context-dependent. MRI could act either as an inhibitor or promoter of NHEJ in a cell-cycle-dependent manner (17, 18). MRI also shares function with XLF, yet has distinct roles, with the combined loss of the two causing pronounced DNA repair defects and severe immunodeficiency due to disrupted V(D)J recombination (13, 19, 20). Despite these genetic and functional insights, the molecular determinants of MRI function remain elusive.

MRI exists in three isoforms: MRI1 (157 residues), MRI2 (69 residues), and MRI3 (102 residues). MRI2 shares the first 46 residues with MRI1, whereas MRI3 encompasses residues 56–157 of MRI1. MRI1 contains two KBMs—an N-terminal APLF-like motif (A-KBM) and a C-terminal XLF-like motif (X-KBM) that has been shown to be important for its function (17, 18, 21). Hydrogen–deuterium exchange experiments suggest that MRI is largely disordered and tends to multimerize (18), but the structure and dynamics of MRI residues are not well characterized. Here, we employed liquid-state NMR to dissect MRI1 and MRI2 at residue-level resolution. We show that the A-KBM of MRI folds upon binding to Ku and that a leucine-to-methionine mutation enhances binding affinity to Ku significantly. Moreover, we identify a previously uncharacterized domain of MRI that drives multimerization and find that this region is important for efficient NHEJ in cells.

## RESULTS

### The A-KBM of MRI is a folding upon binding motif

MRI1 and MRI2 share the same first 46 residues and both contain the same A-KBM (**Fig. 1a**). The A-KBM of MRI binds to the Ku80vWA domain of Ku. The structure of the complex between Ku80vWA and a peptide covering the A-KBM binding motif of MRI has been determined by X-ray crystallography (22). Here, we examine this interaction in the context of the full-length protein MRI. Since MRI1 is prone to aggregation (18), we first used MRI2 to characterize the A-KBM.

**Figure 1.**
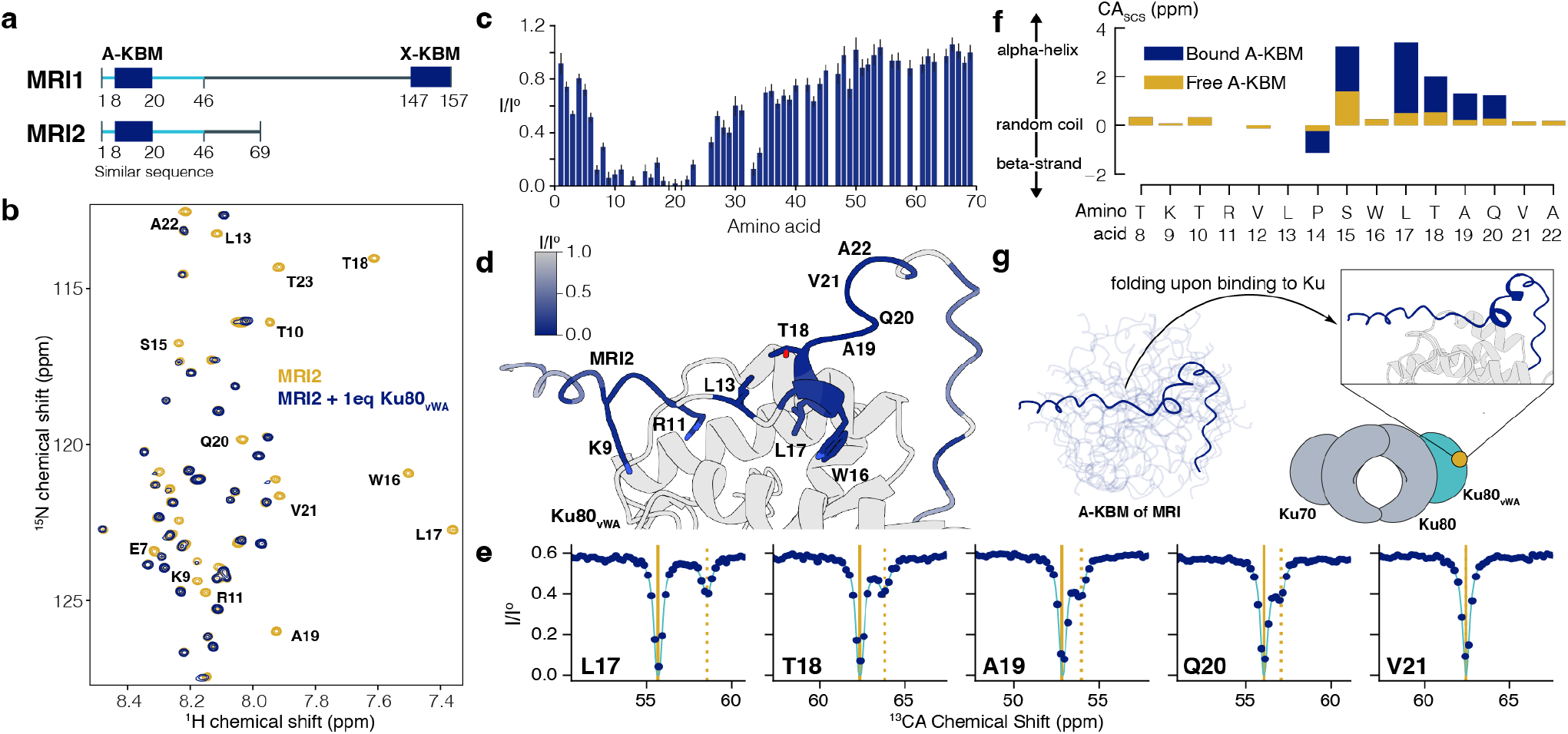
The A-KBM of MRI is folding upon binding to the Ku80_vWA_ domain. (**a**) Domain organization of MRI1 and MRI2 with the indicated APLF-like (A-KBM) and XLF-like (X-KBM) Ku-binding motifs. The region with similar sequence is highlighted in cyan. (**b**) Overlay of ^1^H-^15^N heteronuclear single quantum coherence (HSQC) spectra of 50 µM ^15^N MRI2 alone (yellow) and in the presence of one equivalent of Ku80_vWA_ (blue). (**c**) Peak intensity ratios from ^1^H-^15^N HSQC spectra of ^15^N MRI2 alone (I^o^) and after adding one equivalent of Ku80_vWA_ (I). (**d**) Mapping of the peak intensity ratios in (**c**) on the MRI2-Ku80^vWA^ Alphafold complex model as a color gradient (I/I^0^=0: blue, I/I^0^=1: gray, unassigned residues: white). (**e**) ^13^CA chemical-exchange saturation transfer (CEST) experiment of ^13^C-^15^N MRI2 in the presence of 5% of Ku80_vWA_ domain measured at 293 K on a 950 MHz spectrometer. The solid vertical bar represents the fitted chemical shift of the free state, whereas the dashed bar indicates the fitted chemical shift of the bound form. The ensemble of CEST profiles is given in **Fig. S3**. (**f**) ^13^C_CA_ secondary chemical shifts (CA_SCS_) of residues in the A-KBM of MRI in the free form (yellow) or bound form (blue), fitted from the CEST profiles shown in panel **e** and Fig. **S3**. (**g**) Cartoon illustration of the process of A-KBM folding into a short alpha helix upon binding to the Ku80^vWA^ domain (highlighted in cyan and yellow).

In the ^1^H-^15^N HSQC spectrum of MRI2, the proton chemical shifts are mostly restricted between 7.5 and 8.5 ppm, a signature of disordered proteins. Using a series of 3D NMR experiments, we unambiguously assigned the backbone resonances of all non-proline residues in MRI2 (**Fig. S1a**). We measured ^15^N longitudinal (R_1_) and transverse (R_2_) relaxation rates, as well as heteronuclear nuclear Overhauser effects (hNOE), showing that MRI2 is highly dynamic, with the exceptions of residues around tryptophan 16 in the KBM (**Fig. S1b-d**). The secondary structural propensity calculated from the chemical shifts of the CA and CB atoms shows that the protein mostly exists in a random coil conformational ensemble (**Fig. S1e**). Taken together, these results demonstrate that MRI2 is an intrinsically disordered protein.

Next, we titrated ^15^N-labeled MRI2 with the Ku80_vWA_ domain. The Ku80_vWA_ domain is used instead of the Ku70/Ku80 complex because it can be expressed conveniently in *E. coli*, yielding a high enough quantity for NMR experiments. The addition of one equivalent of Ku80_vWA_ led to the disappearance of several peaks corresponding to the residues in the KBM in the spectra of MRI2. This complete drop in intensity likely comes from a combination of exchange broadening, due to binding occurring in the slow to intermediate exchange regimes and the increase in the reorientational correlation time experienced by MRI2 upon Ku80_vWA_ binding (**Fig. 1b-d**). Interestingly, the residues at the C-terminus of the previously defined KBMs also experience a loss in peak intensity, suggesting that the interactions extends beyond the shorter peptides used in previous studies (22–24).

To gain further insight into the structure and dynamics of the A-KBM of MRI2 upon binding to the Ku80_vWA_ domain, we measured the ^15^N chemical-exchange saturation transfer (CEST) profile of MRI2 in the presence of 10% Ku80_vWA_ (**Fig. S2**). All residues in the A-KBM motif exhibit a ^15^N-CEST profile with two dips, with the minor dip corresponding to the chemical shift of the bound form. Residues 19-22 also show a two-dip chemical exchange profile, suggesting that these residues also undergo structural changes upon binding to the Ku80_vWA_ domain. Fitting the CEST profiles with a two-state model (free and bound states) of all residues together yields an exchange rate of 49.4 *±* 2.4 s^-1^ (**Fig. S2**).

To further characterize the structure of MRI2 in its bound form with Ku80_vWA_, ^13^CO, ^13^CA, and ^13^CB CEST experiments were conducted using ^15^N/^13^C uniformly labeled MRI2 in the presence of a 5% molecular ratio of unlabeled Ku80_vWA_ **(Fig. 1e, S3**). While all residues in the KBM of MRI2 exhibit two dips in the ^15^N CEST experiments, only a few residues show clear minor dips in the ^13^CO, ^13^CA, and ^13^CB CEST experiments. Since carbon chemical shifts are more sensitive to changes in the conformational ensemble, this suggests that the alterations in the secondary structure of MRI2 upon binding to Ku80_vWA_ are not homogeneous. The residues T10, V21, and A22 do not display any minor dips in the three ^13^C CEST experiments, while V12 shows a minor dip only in the ^13^CO CEST experiment (**Fig. 1e, S3**). All residues from S15 to Q20 exhibit a distinct minor dip in the ^13^CO and ^13^CA CEST experiments, except for residue W16. In contrast, residues W16 and T18 show a minor dip in the ^13^CB CEST experiment (**Fig. S3**).

All the CEST profiles with two dips are fitted into a two-state model to extract the chemical shifts of the bound form. The fitted ^13^CA chemical shifts of MRI2 in the bound state were used to calculate the secondary structure propensity (**Fig. 1f**), as the CA chemical shift is the most sensitive to changes in secondary structure. The large positive ^13^CA secondary shift values observed for residues 15-20 in the bound state (25, 26) strongly suggest that these residues form an *α*-helix in the complex, which has not been fully captured in the previous X-ray structure (22). Furthermore, residues 9-14 of the KBM were observed to be in conformation more extended than an α-helix in the complex with Ku, which is consistent with the crystal structures of MRI2_KBM_ and Ku80_vWA_ solved previously (22).

The fitted chemical shifts for the bound form of MRI2 from the ^15^N, ^13^CO, ^13^CA, and ^13^CB experiments were used to predict backbone dihedral angles using TalosN (27). NMR-derived dihedral angles were compared with the X-ray structure and AlphaFold 2 models of the A-KBM in complex with Ku80_vWA_ (**Fig. S4a-b**). The *ψ* and *ϕ* angles derived from the CEST experiments agree well with the backbone conformation of residues modeled in the X-ray structure, with only slight deviations for the first and last residues (**Fig. S4b**). All five models from AlphaFold 2 align well with the calculated dihedral angles from the CEST experiments, including for residues 19-21 that were not modeled in the X-ray structures, with only a slight deviation at residue 21 (**Fig. S4a**).

Taken together, we show that MRI2 is a fully disordered protein, its A-KBM folds upon binding to Ku80_vWA_ (**Fig. 1g**) on the millisecond time scale. The structure of the A-KBM motif in the complex with Ku80_vWA_ shows that residues 9-14 are in a somewhat extended conformation, while residues 15 to 20 form an *α*-helix upon binding.

### L17M mutation in the A-KBM of MRI increases the Ku-binding affinity

The predicted structure and sequence alignment of MRI_KBM_ and APLF_KBM_ reveal that the two structures are expected to be very similar, particularly in the conserved residues (**Fig. 2a-b**), yet APLF_KBM_ has an affinity for Ku80_vWA_ that is two orders of magnitude stronger than MRI_KBM_ (22, 23). Interestingly, sequence alignment of MRI across different species shows that position 17, a leucine in human MRI, is a methionine (M17) in a wide majority of species (**Fig. 2c, S5**). This M17 is also present in the corresponding position in the APLF_KBM_ (**Fig. 2a**). We hypothesized that this position may be crucial for the difference in affinity between KBM_APLF_ and KBM_MRI_.

**Figure 2.**
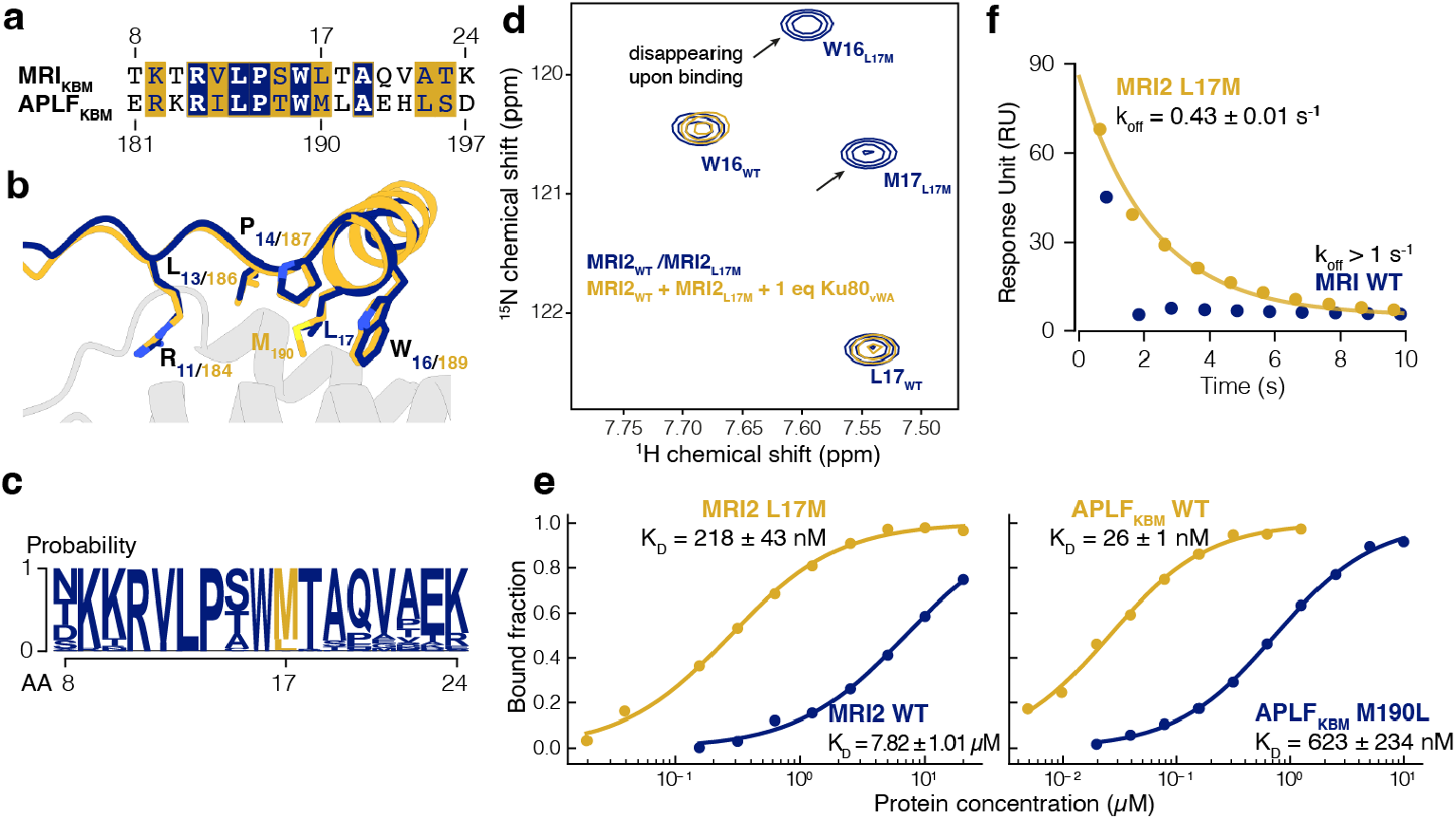
L17M mutation in the A-KBM of MRI increases the affinity for Ku binding. (**a**) Sequence alignment of the residues in the A-KBM of MRI and APLF, strictly conserved residues are highlighted in bold and blue, while regions with conserved amino-acid properties are highlighted in yellow boxes. (**b**) Structural models of Ku80_vWA_ (gray) in complex with the A-KBM of MRI (blue) or APLF (yellow), predicted with AlphaFold 2 multimer (28, 29). (**c**) Logo plot showing the probability of the residues in the A-KBM of MRI from the sequence alignment of MRI2 across different species. (**d**) MRI2 L17M out-competes MRI2 WT in binding to Ku, overlay of ^1^H-^15^N HSQC spectra of 25 µM ^15^N MRI2 WT or MRI2 L17M in isolation (blue) and the mixture of 25 µM ^15^N MRI2 WT and MRI2 L17M with 1 equivalent of the Ku80_vWA_ domain (yellow). (**e**) The bound fraction of MRI2 WT (yellow) or L17M mutation (blue) and APLF_KBM_ WT (yellow) or M190L mutation (blue) was measured by the response at steady states from a single representative surface plasmon resonance (SPR) experiment. The dissociation constant (K_D_) values are displayed as mean ± standard deviation, fitted from n = 9 independent experiments. (**f**) The response in the dissociation phase from a single representative SPR experiment. The value of dissociation rate (k_off_) of MRI2 L17M is displayed as mean ± standard deviation, fitted from n = 3 independent experiments. For MRI2, the k_off_ exceeds the limit of detection of the instrument (k_off_ > 1 s^-1^).

To investigate the role of this leucine to methionine mutation, we tested a series of mutations and constructs of KBM in their interaction with Ku80_vWA_. First, we utilized NMR to observe mixtures comprising ^15^N MRI2_WT_ or MRI2_L17M_ with one equivalent of Ku80_vWA_ (see **Fig. 2d**). Our findings revealed that MRI2 L17M can completely displace MRI2 WT from Ku80_vWA_, as only the signal from MRI2 L17M disappears, while the signal from MRI2 WT remains after the addition of one equivalent of Ku80_vWA_.

Next, we employed surface plasmon resonance (SPR) experiments to quantitatively measure the changes in affinity (through the determination of the dissociation constants K_D_). The L17M mutation in KBM_MRI_ resulted in an increase in affinity by more than 30-fold, reducing the dissociation constant K_D_ from 7.82 µM to 218 nM (**Fig. 2e**). Conversely, introducing the reverse mutation from methionine to leucine (M190L) in KBM_APLF_ reduced the affinity of APLF_KBM_ 24-fold, increasing the dissociation constant K_D_ from 26 nM to 623 nM (**Fig. 2e**).

Comparing the dissociation rates of the interaction between Ku80_vWA_ with MRI2 WT or L17M mutations (**Fig. 2f**), we observed that the increase in affinity is primarily due to a decrease in the dissociation rate. This is likely due to the stabilization of the complex through the methionine mutation, which introduces a slightly bulkier hydrophobic side chain which might better fits at the interface in the complex. Taken together, we conclude that the residue methionine-17 in the KBM of MRI, highly conserved across different species, but not in humans, can significantly enhance the affinity of MRI for Ku.

### Identifying the multimerization domain of MRI1

We used NMR to characterize the first isoform of MRI (MRI1), which contains both the A-KBM and X-KBM motifs. MRI1 is unstable and has been previously shown to form oligomers and aggregates (18). To obtain stable samples of MRI1, the protein was expressed in *E. coli* and purified in buffers containing 3 M urea. The ^1^H-^15^N HSQC spectrum of MRI1 in 3 M urea indicates that the protein is mostly disordered in this condition, with ^1^H chemical shifts primarily in the range of 7.5 to 8.6 ppm. We successfully assigned approximately 90% of the non-proline residues of MRI1. Subsequently, we performed a step-wise dilution of urea from 3 M to 1 M and monitoring the peak intensity (**Fig. S6a**). Upon dilution, the peak intensity significantly decreased across the protein, with a slightly more pronounced reduction for residues 49-81 (**Fig. S6b-c**). We hypothesize that this region may be more structured and prone to aggregation. Indeed, the prediction of disorder propensity further suggests that this domain is likely to be structured (**Fig. 3a**). Additionally, the sequence alignment of MRI1 indicates that the region between residues 49-81 exhibits a high level of conservation (**Fig. 3a**).

**Figure 3.**
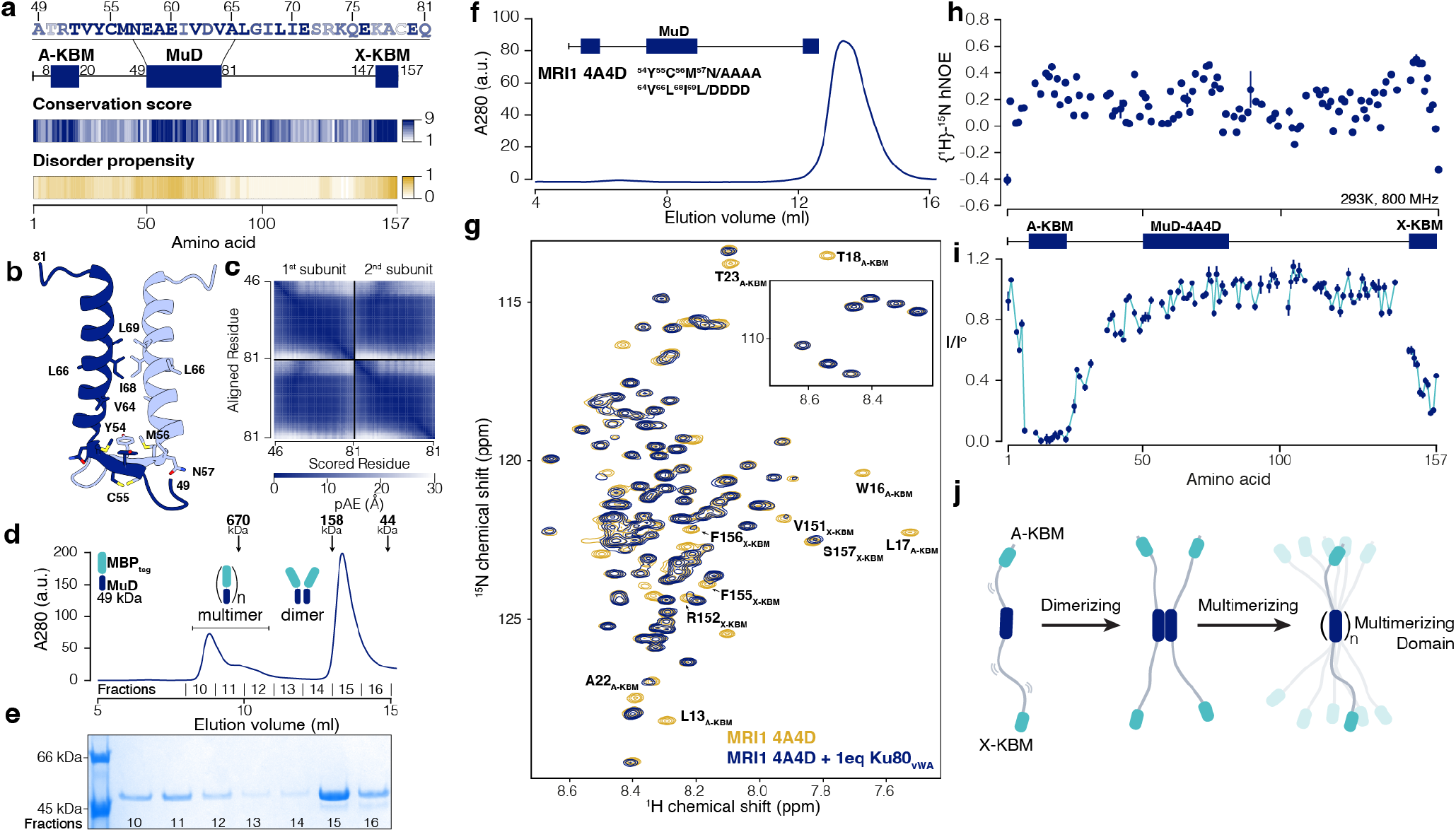
Identifying the multimerization domain of MRI1. (**a**) The conservation score (1: least, 9: most conserved) extracted from sequence alignment of MRI1 across different species by Consurf (30), combined with the disorder propensity (0: disordered, 1: folded) predicted by IUPred2A (31), define the residues between 49-81 as the potential multimerization domain (MuD). (**b**) The structural model and (**c**) the predicted alignment error (pAE) of the MuD dimer predicted with AlphaFold 2 multimer (29). (**d**) The elution profile from size-exclusion chromatography (SEC) of the MuD domain of MRI1 (blue) with the MBP tag (cyan). The elution times for standard proteins with molecular weights of 670, 158, and 44 kDa are indicated with dotted lines. (**e**) SDS-PAGE gel from the indicated elution fractions collected from (**d**). (**f**) The elution profile from SEC of the MRI1 4A4D mutant, combining ^54^YCMN^57^/AAAA and ^64^V^66^L^68^I^69^L/DDDD mutations. (**g**) Overlay of ^1^H-^15^N HSQC spectra of 50 µM ^15^N MRI1 4A4D alone (yellow) and in the presence of one equivalent of the Ku80_vWA_ domain (blue). (**h**) Heteronuclear [^1^H]-^15^N Overhauser effects (hNOE) for residues of MRI1 4A4D mutant are shown. The spectra were recorded at 293 K on an 800 MHz spectrometer. (**i**) Peak intensity ratios from ^1^H-^15^N HSQC spectra of ^15^N spectra of MRI1 4A4D alone (I^o^) and after adding one equivalent of Ku80_vWA_ (I), plotted with the structural organization of MRI1 for comparison. (**j**) Cartoon showing the multimerization process of MRI1 through the multimerizing domain MuD. The A-KBM and X-KBM remain flexible.

We further characterized the domain comprising residues 49-81 of MRI1 (**Fig. 3b-e**). For that purpose, we expressed a fusion protein containing residues 49-81 of MRI1 attached to a N-terminal MBP tag. Gel filtration analysis of the fusion protein indicated the formation of dimers and multimers (**Fig. 3d-e**). AlphaFold 2 predicted the structure of the 49-81 domain as a dimer, with moderate confidence (interface predicted template modelling score, ipTM of 0.42) and a low predicted alignment error (**Fig. 3b-c**). The dimer model shows the packing of two alpha helices together from residues 58 to 81, while residues 53-57 form a beta strand that can pair with the other subunit. Taken together, we conclude that the region encompassing residues 49-81 drives the formation of MRI1 dimers and multimers, which we refer to as the multimerizing domain (MuD).

Based on the structural model of the MuD dimer from AlphaFold 2, we introduced several mutations to disrupt the dimer, including four alanine mutations in the beta strand (^54^YCMN^57^/AAAA) and four aspartic acid mutations in the alpha-helix region (^64^V^66^L^68^I^69^L/DDDD), collectively termed the 4A4D mutation (MRI1 4A4D). These mutations render MRI1 completely soluble even at high concentrations and result in a single peak elution from the gel filtration column (**Fig. 3f**), indicating the protein solution is monodisperse. We assigned backbone chemical shift to ca. 92% of the non-proline residues of this construct. The ^1^H-^15^N HSQC spectrum, along with ^15^N relaxation data including hNOE, longitudinal and transverse relaxation rates all show that the protein is mostly disordered (**Fig. 3g-h, S7**), but slightly less dynamic than MRI2.

Since MRI1 contains both A-KBM and X-KBM at N-terminal and C-terminal extremities, respectively, we next characterized the interaction between MRI1 and Ku. Adding an equimolar amount of the Ku80_vWA_ domain to ^15^N MRI1 4A4D led to the disappearance of the peaks from several residues corresponding to the A-KBM and X-KBM (**Fig. 3g,i**). This also shows that both A-KBM and X-KBM can interact with Ku80_vWA_ simultaneously. In summary, we identify a structural region (residues 49-81) in MRI1 that promotes oligomer formation, while the rest of the protein is highly flexible with the A-KBM and X-KBM at the two extremities are both capable of binding to Ku80_vWA_.

### Multimerization of MRI1 is essential for stimulating DSB repair by NHEJ *in vivo*

With insights from the characterization of molecular properties of MRI1, we designed various mutants of MRI1 and evaluated their impact on NHEJ activity in cellular models (**Fig. 4a, S8**). We focused solely on MRI1, as this isoform has been shown to exhibit a phenotype upon knockout, whereas an MRI2 knockout does not display a significant phenotype (13, 17).

**Figure 4.**
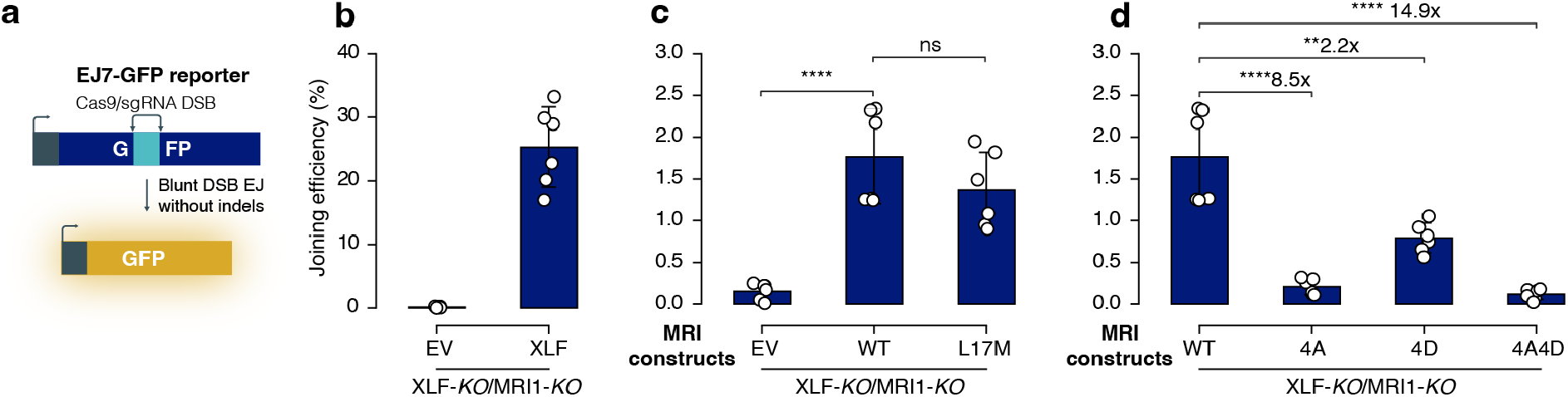
Multimerization of MRI1 is essential for stimulating DSB repair by NHEJ *in vivo*. (**a**) The EJ7-GFP reporter (not to scale) for No Indel EJ between two Cas9/sgRNA-induced DSBs, integrated into XLF-*KO*/MRI1-*KO* HEK293 cells. (**b-d**) Joining efficiency of XLF, wild-type (WT) MRI1, and different mutants (4A: MRI1 ^54^YCMN^57^/AAAA, 4D: MRI1 ^64^V^66^L^68^I^69^L/DDDD, 4A4D: MRI1: MRI1 ^54^YCMN^57^/AAAA-^64^V^66^L^68^I^69^L/DDDD) in the XLF-MRI1 double knockout cells. The normalized joining efficiency was assessed by GFP frequencies, accounting for transfection efficiency. n = 6 biologically independent transfections. Statistical significance was determined by an unpaired two-tailed t-test: ****P<0.0001, ***P<0.001, and **P<0.01, ns: not significant, EV: empty vector.

We utilized the EJ7-GFP reporter to assess end joining between the distal ends of two Cas9-mediated blunt chromosomal DSBs, ensuring that no indel mutations occurred at the edges of the DSB (**Fig. 4a**). The significance of blunt DSB end joining without indels lies in the fact that this type of repair is not stabilized by an annealing intermediate and has been demonstrated to rely on NHEJ (32). This no-indel end joining results in the restoration of a GFP+ cassette, which is quantified using flow cytometry. To disentangle the functional overlap between XLF and MRI1, we knocked out both proteins and subsequently complemented the cells with either XLF or different variants of MRI1.

A double knockout of MRI1 and XLF impairs total NHEJ activity, with joining efficiency nearly at zero (**Fig. 4b**). When XLF is added to the double knockout, NHEJ activity is partly rescued, as the joining efficiency increases to 20-30% (**Fig. 4b**). In contrast, complementing the cells with MRI1 alone restores the joining efficiency to 1-2.5% but significantly more than the empty vector (**Fig. 4b-c**). Having identified the L17M mutation that can increase the affinity of A-KBM for Ku by over 30 times, we designed an L17M mutation in MRI1. Despite the significant increase in affinity, MRI1 with the L17M mutation did not statistically change the joining efficiency (**Fig. 4c**).

We tested the impact of multimerization of MRI1 on NHEJ activity. We designed three MRI1 constructs with mutations that disrupt multimerization: MRI1 4A, which contains four mutations in the putative beta-strand (^54^YCMN^57^/AAAA mutations), MRI1 4D, which has mutations in the putative alpha-helix (^64^V^66^L^68^I^69^L/DDDD mutation) and MRI1 4A4D, which includes both sets of mutations (**Fig. 3b, f**).

All three constructs exhibited a marked reduction in NHEJ activity compared to the wild type (**Fig. 4d**). The introduction of the 4A or 4A4D mutations nearly completely disrupted NHEJ activity, reducing it to levels comparable to that of the empty vector. In contrast, the 4D mutations in the alpha-helix resulted in a lesser reduction in activity compared to the other two constructs. This indicates that the 4A mutations in the beta-strand are more disruptive to the formation of the oligomer and suggest that the beta-strand contributes more significantly to oligomer formation than the alpha-helix. Taken together, these findings indicate that the multimerization of MRI1 by the MuD domain is essential for stimulating NHEJ activity.

## Discussion

In recent years, the molecular mechanism of NHEJ has been characterized extensively, revealing the intricate complexity of NHEJ across different scales and dynamics (3, 5–9). However, several critical aspects remain to be explored, particularly the role of auxiliary factors. The structural characterization of these auxiliary NHEJ factors presents additional challenges, as many contain multiple intrinsically disordered regions (IDRs) that are mostly invisible in X-ray crystallography and challenging to interpret in cryoEM electron densities. Furthermore, functional redundancy among auxiliary factors complicates their characterization, as individual protein knockdowns often produce only modest phenotypic effects due to overlapping roles (12, 13, 33). Despite these experimental challenges, recent research has begun to shed light on the contributions of auxiliary factors to the NHEJ pathway, as well as the role of the IDRs (10–12).

The affinity of IDRs for partner proteins can be fine-tuned depending on the specific context (34). In this study, we demonstrate that a single mutation from leucine to methionine in the A-KBM significantly enhances the affinity of KBM for Ku. This mutation may have evolved to accommodate different functions across various proteins. For instance, APLF exhibits a high affinity for binding, which is necessary to stabilize the NHEJ synapsis (11), while the same mutation may have evolved in different species to optimize the activity of DNA repair machinery. We found that the M17 residue is more predominantly conserved than L17 across species, with the exception of humans and closely related primates. The methionine-to-leucine mutation in MRI_KBM_ decreases the affinity of MRI_KBM_ for Ku by 30-fold. Given that the concentration of Ku is significantly higher in human cells compared to other species, such as mice, cows, and whales (35), it is plausible that the MRI_KBM_ has evolved as an adaptation to balance the elevated levels of Ku in those species. Since dysregulation of MRI can lead to chromosome fusion (17), it is crucial that the affinity of MRI for Ku is fine-tuned to prevent genomic instability resulting from hyperactive NHEJ activity. Although we show that in the EJ7 assay, the L17M mutation does not significantly change NHEJ activity, it is important to note that this assay is an endpoint measurement. We do not rule out the possibility that this mutation may alter the kinetics of the repair process, which warrants further investigation.

The kinetics of interactions and dynamics of IDRs are often advantageous compared to folded proteins, particularly in protein-protein interactions. With comparable affinity, IDRs can associate and dissociate with their binding partners more rapidly than folded proteins (36). Our findings indicate that the A-KBM of MRI can remain predominantly disordered while folding upon binding to Ku (**Fig. 1**). Coupled with a long flanked disordered region, the A-KBM and X-KBM can bind to Ku over a larger radius, akin to a “fly-catch mechanism” (37), rendering MRI as an effective adaptor for NHEJ synapsis as shown by previous studies (13, 17, 18, 38). We also identified the domain that mediates MRI multimerization (MuD) and demonstrate that the multimerization is important for promoting NHEJ activity. We further propose that multimerization of MRI1 through the MuD, coupled with the KBMs at both termini, enables multivalent interaction of MRI with Ku and thereby stimulates NHEJ activity (**Fig. 5**), similar to the roles described for XLF and XRCC4 (10). However, unlike XLF and XRCC4, MRI1 does not form condensates *in vitro* (18, 38) or in cells (17).

**Figure 5.**
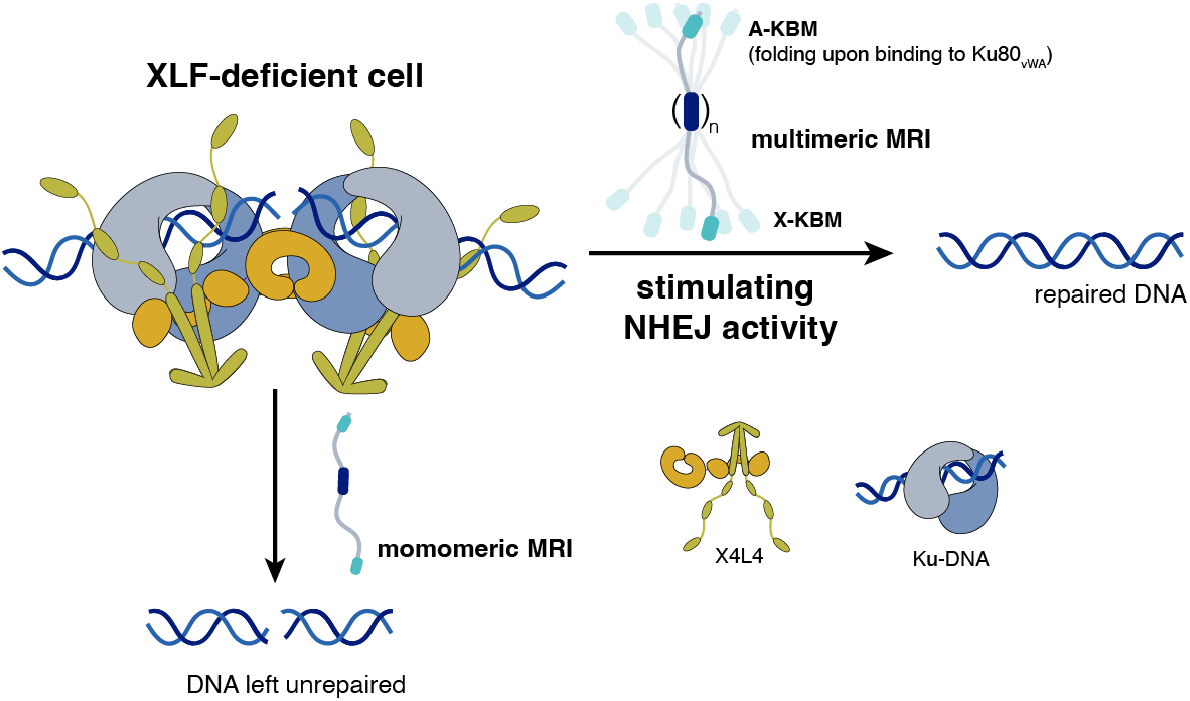
Cartoon illustrating the multimerization of MRI1 leading to the stimulation of NHEJ in DNA double-strand break repair in the context of XLF-deficient cell.

How multimeric, but not monomeric, MRI promotes NHEJ, and at which stage of synapsis this activity becomes important, remain to be determined. The mutants characterized in this study provide a useful framework for addressing these questions in future work. The functional significance of MRI multimerization may also extend beyond its role in NHEJ. In addition to NHEJ factors, MRI has been reported to associate with proteins involved in the broader DNA damage response and homologous recombination repair (18). Moreover, both MRI and Ku have recently been linked to functions in RNA splicing (38, 39). By presenting multiple Ku-binding sites, it is possible that multimeric MRI facilitates Ku recruitment to sites of DNA damage or RNA splicing.

In summary, we have explored the dynamics of two MRI isoforms, described the binding of the A-KBM to Ku in the context of full-length MRI and characterized the functional significance of MuD domain of MRI1, which provide the molecular basis for further investigation to gain a deeper understanding of the role of MRI in regulating NHEJ activity. Our work emphasizes the important role of IDRs within the NHEJ mechanism and may have potential implications for both therapeutic and biotechnological applications.

## METHODS

### Plasmid constructs

Wild-type MRI1 (Q9BWK5-1), MRI2 (Q9BWK5-4), and the indicated mutants were fused with MBP, an 8xHis tag, and a TEV cleavage site at the N-terminus, and then subcloned into the pMAL-c5X plasmid. All plasmids were synthesized and sequenced by Gene Universal Company. For wild-type MRI1, the protein was also subcloned into pET-28b with an 8xHis tag and a TEV cleavage site at the N-terminus.

### Bioinformatic analysis

Sequence alignments of MRI1 and MRI2, as well as phylogenetic trees, were generated using the ConSurf web server with default parameters (30). Intrinsic disorder propensity prediction for MRI1 was performed using IUPred2A with default parameters (31). Structural predictions of wild-type MRI2 and its mutants in complex with the Ku_vWA_ domain were generated using AlphaFold2 that is implemented in ColabFold, with default parameters and five different models (29, 40).

### Protein expression and purification

**MRI1 and MRI2:** The plasmids containing WT and mutated MRI2, as well as mutated MRI1, were used to transform into *E. Coli* Novagen BL21 Rosetta^TM^ (DE3) bacteria. The bacteria were grown at 37 °C in lysogeny broth (LB) containing 75 µg/ml ampicillin and 25 µg/ml chloramphenicol until reaching an optical density at 600 nm between 0.6 and 0.8. Protein expression was subsequently induced with 500 µM isopropyl β-D-1-thiogalactopyranoside (IPTG) and cells were incubated overnight at 16 °C. The pellets were collected by centrifugation at 11000×g at 4 °C for 30 minutes and sonicated in lysis buffer (50 mM Tris-HCl pH 7.5, 1000 mM NaCl, 2 mM DTT, 1 mM EDTA, 100 mM arginine and glutamic acid, 5% (v/v) glycerol). The supernatant was obtained by centrifuging at 11000×g for 45 minutes at 4 °C and loaded into the prepacked HisTrap^TM^ excel 5ml column preequilibrated with buffer A1 (25 mM TrisHCl pH 7.5, 50 mM NaCl , 2 mM DTT, 1 mM EDTA) at the rate of 1 mL/min and was washed extensively with 20x column volume of buffer B1 (25 mM TrisHCl pH 7.5, 1000 mM NaCl, 2 mM DTT, 1 mM EDTA). The protein was then eluted with 80% buffer A2 (50 mM TrisHCl pH 7.5, 50 mM NaCl, 2 mM DTT, 1 mM EDTA, 300 mM imidazole) + 20% buffer B1. The His tag was cut by adding 1% tobacco etch virus (TEV) protease (w/w) to the samples while dialyzing in buffer A1 overnight. The following day, the protein was passed through the HiTrap S 5 mL column and the proteins were eluted with a salt gradient between 0-60% B1 with a ramp increase of of 1% per minute. After that, the fractions that contained the protein of interest were concentrated and passed through a SEC 16/60 S75 column pre-equilibrated with NMR buffer (20 mM Bis-Tris pH 6.5, 150 mM KCl, 1 mM DTT, 1 mM EDTA). The fractions corresponding to the peaks were identified by electrophoresis in a 15% precast polyacrylamide gel, pooled together, and concentrated using 3 kDa molecular weight cut-off concentrator, flash frozen in liquid nitrogen and stored at −80 °C.

The ^15^N and ^15^N/^13^C uniformly labeled samples were prepared using the Marley method (41) with ^15^NH_4_Cl and ^13^C glucose as the sole sources of nitrogen and carbon nutrients, respectively.

For MRI1, due to the high propensity of aggregation, the protein was purified using the same protocol as mentioned above (keeping the His tag) with the addition of 3 M Urea in all buffers, then passed through a SEC 16/60 S75 column preequilibrated with NMR buffer + 3 M urea.

**Ku:** Ku80/Ku70_SNAP_ and Ku80_vWA_ were expressed and purified as previously described (24) and (10), respectively.

### Cellular GFP reporter NHEJ assay

The HEK293 EJ7-GFP cell line and XLF-KO/MRI-KO cells, and plasmids for the EJ7 reporter assay, and the expression vector for pCAGGS-3xFLAG-MRI-WT were previously described (13). The pCAGGS-3xFLAG-MRI-WT plasmid was used to generate the mutant versions.

For the EJ7-GFP reporter assay, HEK293 cells were plated at 2×105 in a poly-lysine-coated 24-well plate. The following day, cells were transfected with 200ng of sgRNA/Cas9 plasmids (7a and 7b), as well as 50ng of MRI-WT, MRI-L17M, MRI-4A, MRI-4D, MRI-4A4D, or EV control (pCAGGS-BSKX) plasmid. Parallel transfections were tested with 200 ng of GFP expressing plasmid (pCAGGS-NZE-GFP), 200 ng of EV, and 50 ng of MRI-WT, MRI-L17M, MRI-4A, MRI-4D, MRI-4A4D, or EV control plasmid. Each well used 1.8 uL of Lipofectamine 2000 (Thermofisher) and 1.5 mL of antibiotic-free media. Three days after transfection, flow cytometry (ACEA Quanteon, Agilent NovoExpress Version 1.5.0) was used to analyze cells, as described (42).

For immunoblot analysis, transfections were as for the reporter assay, but scaled 4-fold, and replaced the Cas9/sgRNA plasmids with EV. Cells were scraped and lysed in ELB buffer (250 mM NaCl, 5 mM EDTA, 50 mM Hepes, 0.1% (v/v) Ipegal, and Roche protease inhibitor) with sonication (Qsonica, Q800R). Immunoblots were probed with antibodies for Anti-Flag HRP (Sigma A8592) and Tubulin (Sigma T9026). ECL reagent (Amersham Biosciences) was used to develop HRP signals.

### NMR

#### MRI1 and MRI2

Unless otherwise noted, the NMR spectra were obtained at 293 K using a Bruker 18.8 T (800 MHz proton frequency) spectrometer equipped with a room-temperature triple resonance (TXI) probe. The NMR data were processed using NMRPipe (43) and analyzed and visualized with NMRFAM-Sparky (44) or POKY (), as well as matplotlib 3.8, seaborn 0.13, and Python 3.9. The sample buffer contained 20 mM BisTris at pH 6.5, 150 mM KCl, 1 mM EDTA, 1 mM DTT, and 5% D_2_O. To assign MRI1 and MRI2, a series of 3D BEST-TROSY experiments, including HNCA, HN(CO)CA, HNCACB, HN(CO)CACB, HN(CA), and HNCO (45–47), were performed on uniformly labeled ^13^C and ^15^N samples of MRI1 4A4D, and MRI2, with concentrations between 250 and 500 µM. Additionally, for MR1 WT, a 4D HNCOCA spectrum was acquired at 293 K on a Bruker 14.1 T spectrometer using non-uniform sampling.

#### NMR Relaxation Measurement

The ^15^N longitudinal relaxation rates (R_1_) were determined by sampling the decay of of longitudinal polarization with eight to ten delays ranging from 0 to 1.8 s. All ^15^N transverse relaxation rates (R_2_) were measured using a Carr-Purcell-Meiboom-Gill (CPMG) train of ^15^N *π*-pulses interleaved with ^1^H π-pulses (48, 49). Six delays between 4 and 120 milliseconds were utilized to determine the transverse relaxation rates R_2_. The ^1^H-^15^N heteronuclear nuclear Overhauser effects (hNOE) were measured by detecting the steady-state polarization of ^15^N while saturating the protons with a series of π-pulses, comparing the peak intensities to those from an experiment where the ^15^N magnetization had returned to equilibrium (50).

#### NMR Titration Experiments

In protein-protein titration experiments, unlabeled proteins were added to solutions of isotopically labeled proteins with the amount as indicated. Chemical shift perturbation (CSP) values for all residues were calculated using the following equation: 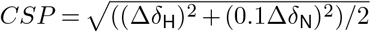, where Δ*δ*_H_ and Δ*δ*_N_ are the changes of chemical shifts for proton and nitrogen-15 respectively.

#### Chemical-exchange NMR experiments

The chemical-exchange saturation transfer (CEST) experiments were acquired at 800 ^1^H MHz (^15^N CEST) or 950 ^1^H MHz (^13^CA, ^13^CB, ^13^C’O CEST) spectrometer with previously published pulse sequences (51–56) on uniformly ^15^N-(^15^N-CEST) and ^15^N,^13^C-(^13^C-CEST) labeled samples of MRI2 (400 *µ*M) with a 5% or 10% molar ratio of unlabeled Ku80_vWA_ as indicated. Relaxation delays (*T*_ex_) were set to 300 ms for all experiments.

For ^15^N CEST, a series of ^15^N DANTE-CEST experiments was conducted using the previously published pulse sequence (55), with CEST spectral width (sw_CEST_) values of 500, and 700 Hz centered at 119.232 ppm, corresponding to effective DANTE rf values of 10 and 20 Hz, respectively.

For ^13^C’O CEST, a pair of ^13^C’O cosine-modulated CEST experiments was performed using the previously published pulse sequence (56), with spectral widths of 1040 or 1200 Hz centered at 175.757 ppm.

For ^13^CA and ^13^CB CEST, a series of 2D ^15^N-^1^H planes were collected in an interleaved manner, with CEST irradiation offsets incremented from 47.74 to 68.67 ppm for ^13^CA CEST and from 16.84 to 47.18 ppm for ^13^CB CEST in steps of 50 Hz for both experiments. For ^13^CB CEST, a separate experiment was conducted with CEST irradiation offsets incremented from 66.85 to 73.76 ppm to focus on threonine residues (54, 57).

### Surface plasmon resonance experiments

Surface plasmon resonance (SPR) experiments were conducted at 25 °C using a Biacore T200 (GE Healthcare Life Sciences) with a Sensor Chip SA Series S (#BR100531, Cytiva). The Ku70-Ku80 complex, with a Snap tag on Ku70, was expressed as previously described (24). SNAP-Biotin (#S9110S, New England Biolabs) was combined with Ku70/Ku80-Snap in a 1:10 ratio and incubated for 2 hours. Subsequently, a buffer exchange was performed with the SPR running buffer, which is similar to the NMR buffer but includes 0.01% Tween 20 (20 mM BisTris, pH 6.5, 150 mM KCl, 1 mM EDTA, 0.5 mM TCEP + 0.01% Tween 20). The flow rate was maintained at 30 µL·min^-1^. Biotinylated Ku was immobilized using the response target mode, with the target set to 2500 response units. A flow cell without immobilized Ku served as a negative control, while the other three flow cells with immobilized Ku acted as active flow cells. Each concentration was injected three times into both the reference and active flow cells. The analyte and subsequent buffer injections were set between 30 and 60 seconds, respectively, at a flow rate of 30 *µ*L·min^-1^. Regeneration was achieved by a pulse injection of 1 M NaCl. Data were collected at a rate of 1 Hz. All sensorgrams were adjusted by reference subtraction. Data analysis was performed using Steady-State Analysis with the Biacore T200 Evaluation software (GE Healthcare) using a 1:1 binding model.

### Statistics and Reproducibility

Statistical significance was assessed using an unpaired two-tailed t-test. The sample sizes selected for all experiments were based on the experiences of our laboratory and other researchers with comparable assays, ensuring both reproducibility and statistical significance.

## Acknowledgements

We thank Alicia Vallet for helping with the NMR experiments. **Funding:** This work was funded by the French ANR (ANR-18-CE29-0003 NANO-DISPRO, ANR-17-CE2-0020 NHEJLIG4, ANR-23-CE11-0039-01 XXL). This work has been supported by the Fondation ARC pour la recherche sur le cancer (grant N°ARCDOC42021120004347) as well as the Equipex contract ANR-10-EQPX-09 (Paris en resonance). Financial support from the IR INFRANALYTICS FR2054 CNRS for conducting the research is gratefully acknowledged. The funders had no role in study design, data collection and analysis, decision to publish or preparation of the manuscript.

## Authors contributions

F.F., M.M., D.D.V. conceived the project and obtained the main funding. D.D.V. prepared reagents and performed the NMR and *in vitro* experiments with the help of S.K., P.P., L.C., G.B.; M.M., K.M., J.M.S. prepared reagents and performed the cellular experiments. D.D.V., M.M. and F.F. wrote the manuscript and all the co-authors edited and approved the final version of the manuscript.

## Data availability

The NMR assignments and relaxation data for MRI1 (4A4D) and MRI2 can be accessed in the Biological Magnetic Resonance Data Bank (BMRB) with the following codes: 53894, 53895. The previously published structures of proteins were retrieved from PDB under the following code: MRI2-Ku_vWA_ (6TYU). All data supporting the findings is provided with the paper in the source file. Other materials generated in this work will be made available upon request.

