## Supplementary figures for "Molecular characterization of MRI/CYREN reveals the Ku binding mode and the role of multimerization in stimulating the activity of NHEJ in DNA repair"

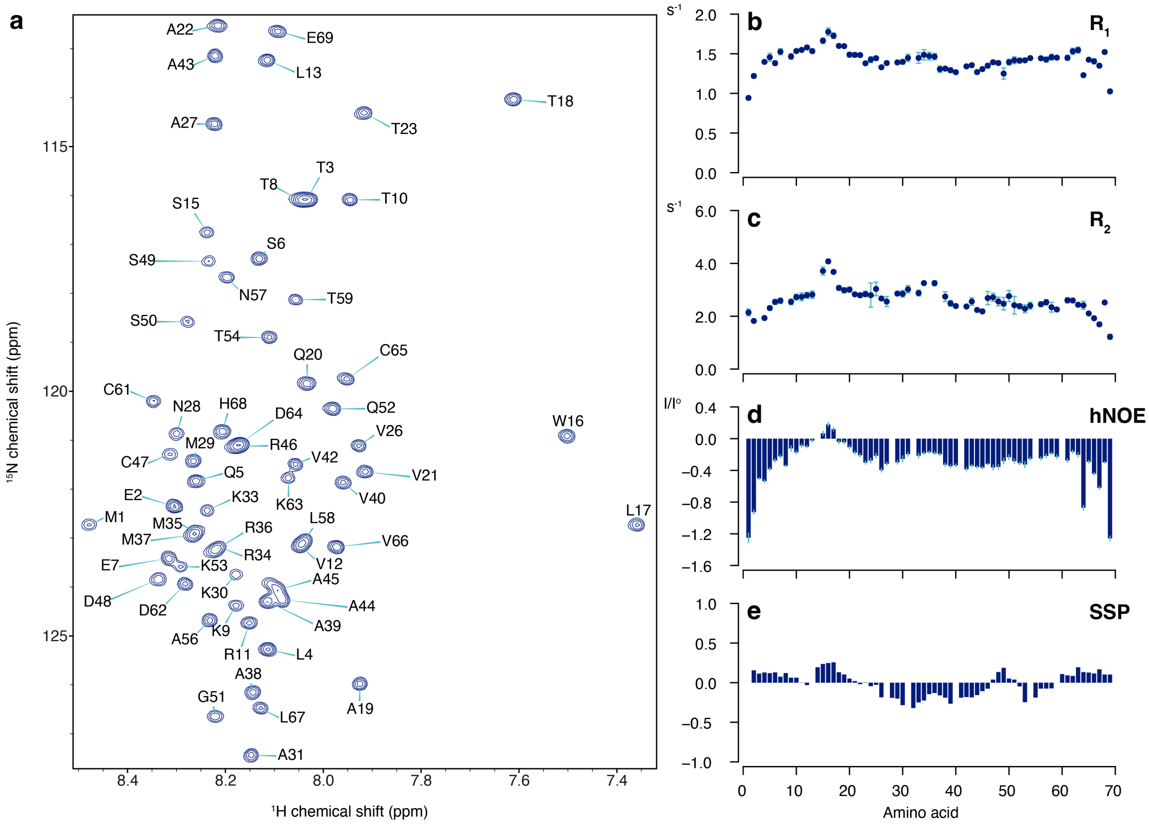


**Figure S1. MRI2 is an intrinsically disordered protein.** (**a**) NMR resonance assignment of the $^{1}H$-$^{15}N$ HSQC spectrum of MRI2, (**b**) $^{15}N$ longitudinal relaxation rate ($R_1$), (**c**) CPMG transverse relaxation rate ($R_2$), (**d**) heteronuclear [$^{1}H$]-$^{15}N$ Overhauser effects (hNOE), (**e**) secondary structure propensity of MRI2 calculated using $C^{\alpha}$ and $C^{\beta}$ chemical shifts. The error bars represent the uncertainty with one standard deviation, estimated either from the covariance matrix (**b-c**) or noise propagation (**d**), with n=1 independent experiment for each spectrum. The experiments were measured on a 600 MHz spectrometer at 293 K.

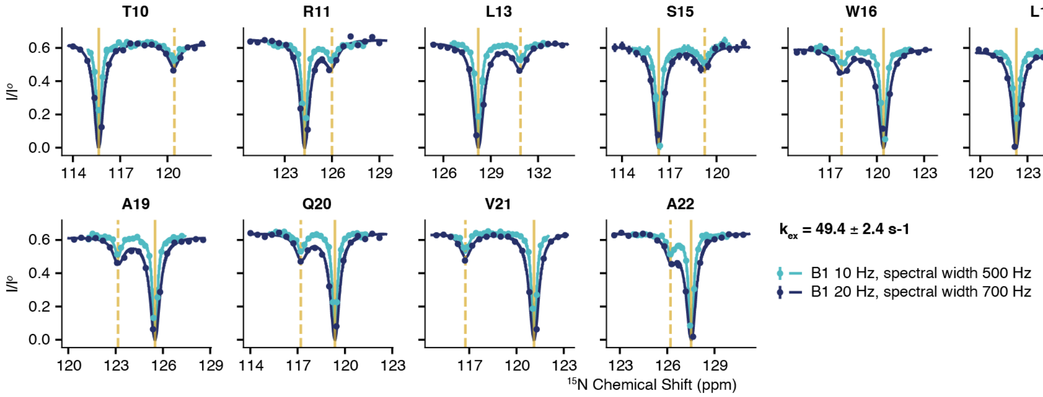


**Figure S2. $^{15}$N DANTE-CEST Experiments of MRI2.** $^{15}$N CEST experiments of MRI2 in the presence of a 10% molar ratio of Ku80$_{vWA}$ were conducted with two effective rf amplitudes: 10 Hz, spectral width 500 Hz (cyan) and 20 Hz, spectral width 700 Hz (blue). The experiments were measured on an 800 MHz spectrometer at 293 K. The solid vertical bar represents the fitted chemical shift of the free state, whereas the dashed bar indicates the fitted chemical shift of the bound form, refer to the supplementary data for the fitted data.

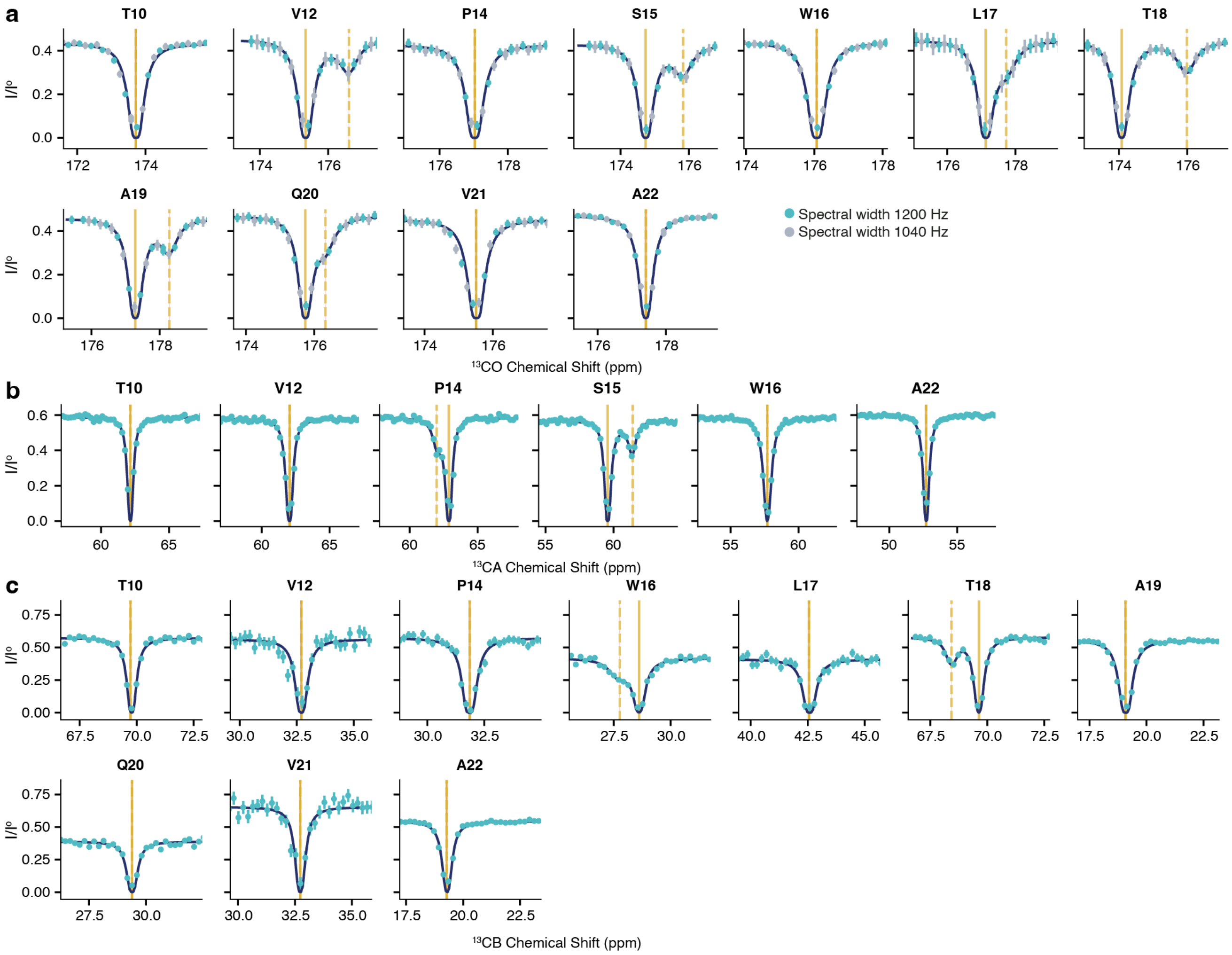


**Figure S3. $^{13}$C CEST experiments of MRI2.** $^{13}$CO (**a**), $^{13}$CA (**b**), and $^{13}$CB (**b**) CEST experiments of $^{15}$N-$^{13}$C MRI2 in the presence of 5% Ku80$_{vWA}$. The experiments were measured on a 950 MHz spectrometer at 293 K. The error bars represent the uncertainty by assuming additive Gaussian noise on the profile. The solid vertical bar represents the fitted chemical shift of the free state, whereas the dashed bar indicates the fitted chemical shift of the bound form, refer to the supplementary data for the fitted data.

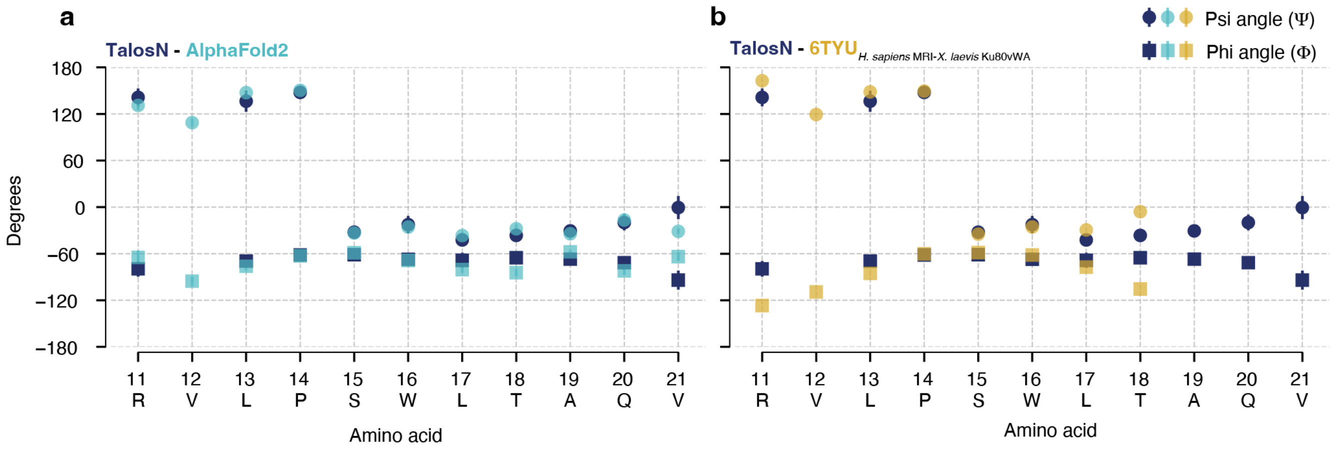


**Figure S4. Backbone dihedral angles of the bound form of MRI2 obtained by chemical-exchange NMR.** The $^{15}N_H$, $^{13}CA$, $^{13}CB$, and $^{13}CO$ chemical shifts of MRI2 in its bound form, extracted from $^{15}N$ and $^{13}C$ CEST experiments, were used to predict the $\psi$ (circle) and $\phi$ (square) angles of MRI2 bound to $Ku80_{vWA}$ using TalosN (1) (dark blue). These values are compared with the angles extracted from AlphaFold structure models (cyan, **a**) and the crystal structure of *H. sapiens* MRI in complex with *X. laevis* $Ku80_{vWA}$ (yellow, **b**), PDBid: 6TYU. The error bars for TalosN values represent the uncertainty of the model, while the error bars for the values extracted from AlphaFold2 structure models indicate the standard deviation calculated from five model outputs of AlphaFold2.

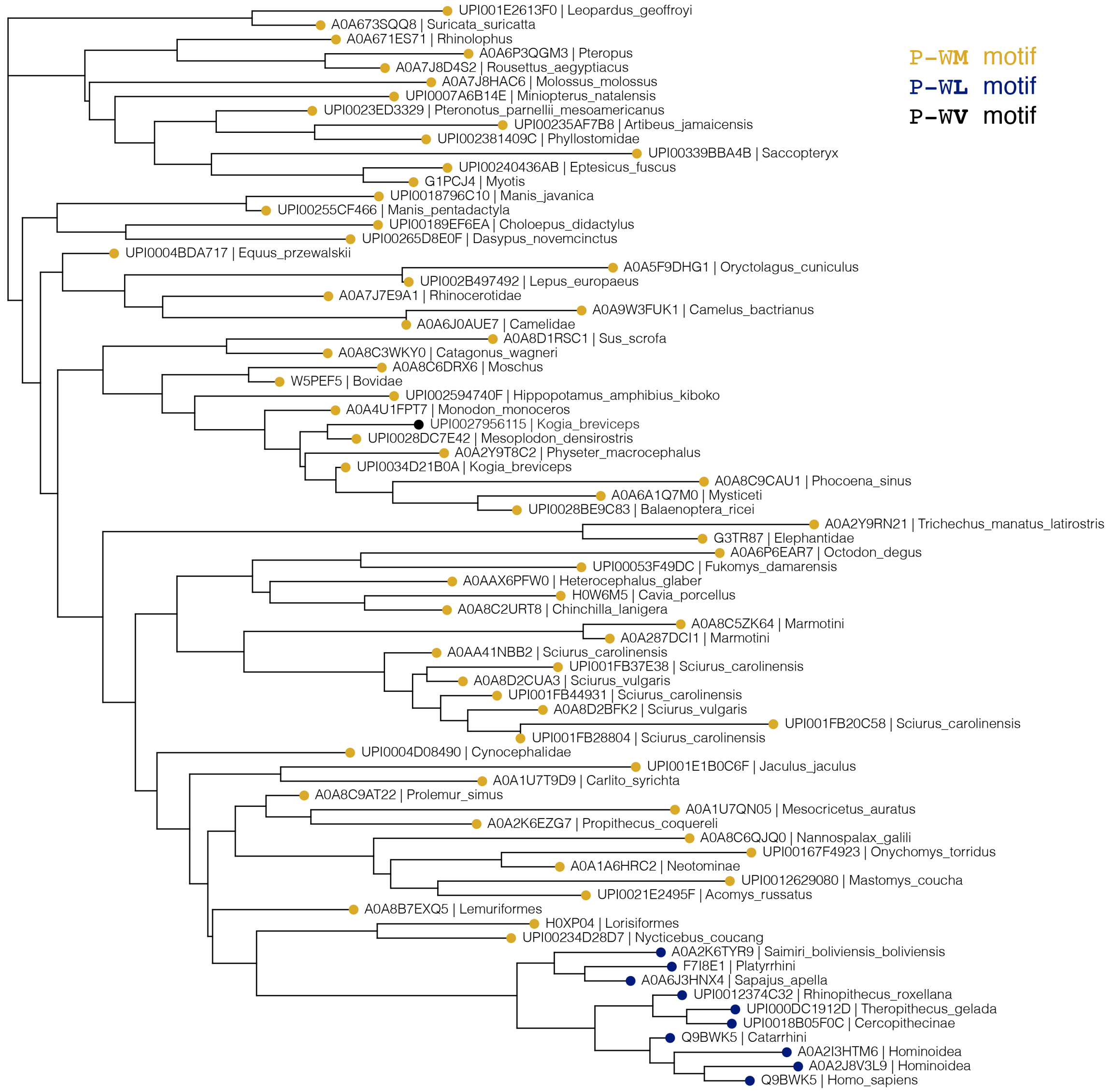


**Figure S5. M17 is more dominantly conserved than L17 in the A-KBM of MRI.** The phylogenetic tree of MRI2 was constructed using 76 unique sequences from the evolutionary conservation analysis conducted with the Consurf web server (2).

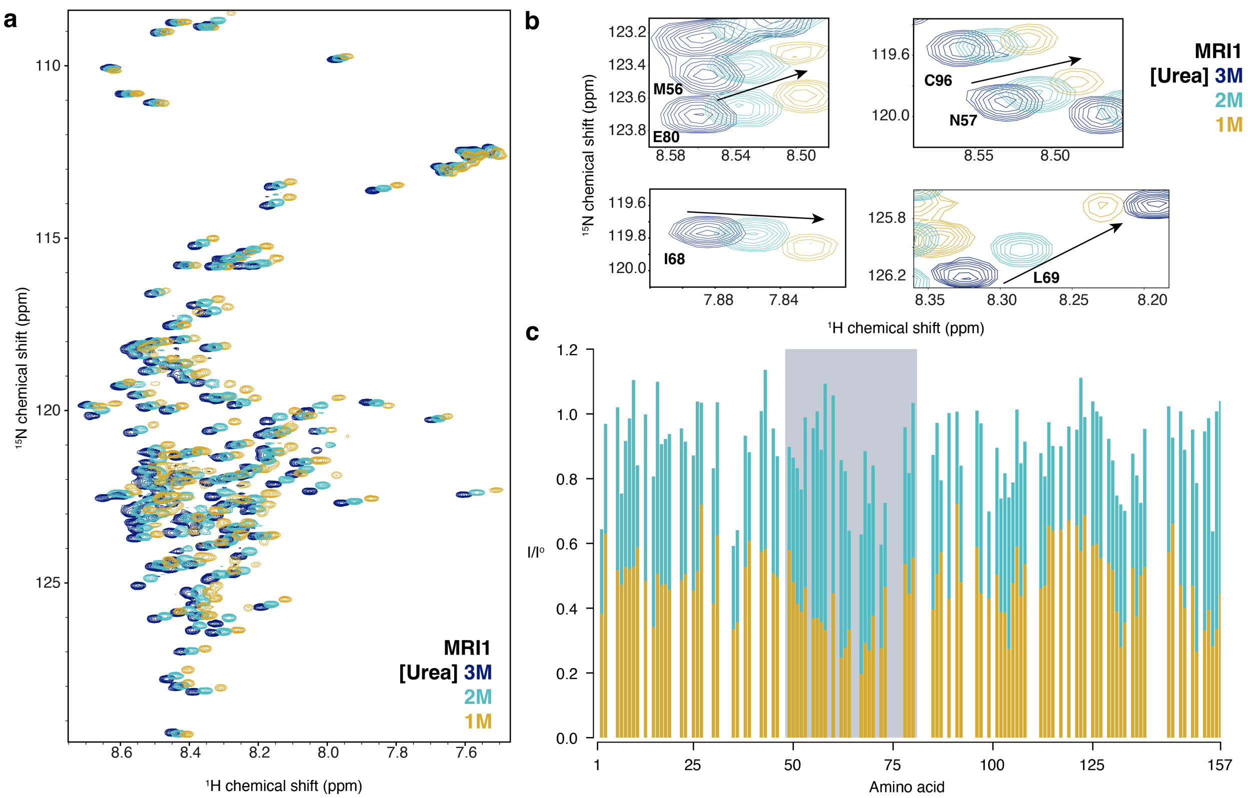


**Figure S6. MRI1 in the presence of different concentrations of Urea** (**a**) Overlay of $^{1}$H-$^{15}$N HSQC spectra of MRI1 in the presence of 3 M (blue), 2 M (cyan) and 1 M (yellow) Urea. (**b**) View of the peaks in a region where the signals of the residues in the multimerization domain of MRI1 appear. (**c**) Peak intensity ratios (I/I$^{O}$) from $^{1}$H-$^{15}$N HSQC spectra of $^{15}$N MRI1 in the presence of 3M Urea (I$^{O}$) and 2 M Urea (I, cyan) or 1 M Urea (I, yellow).

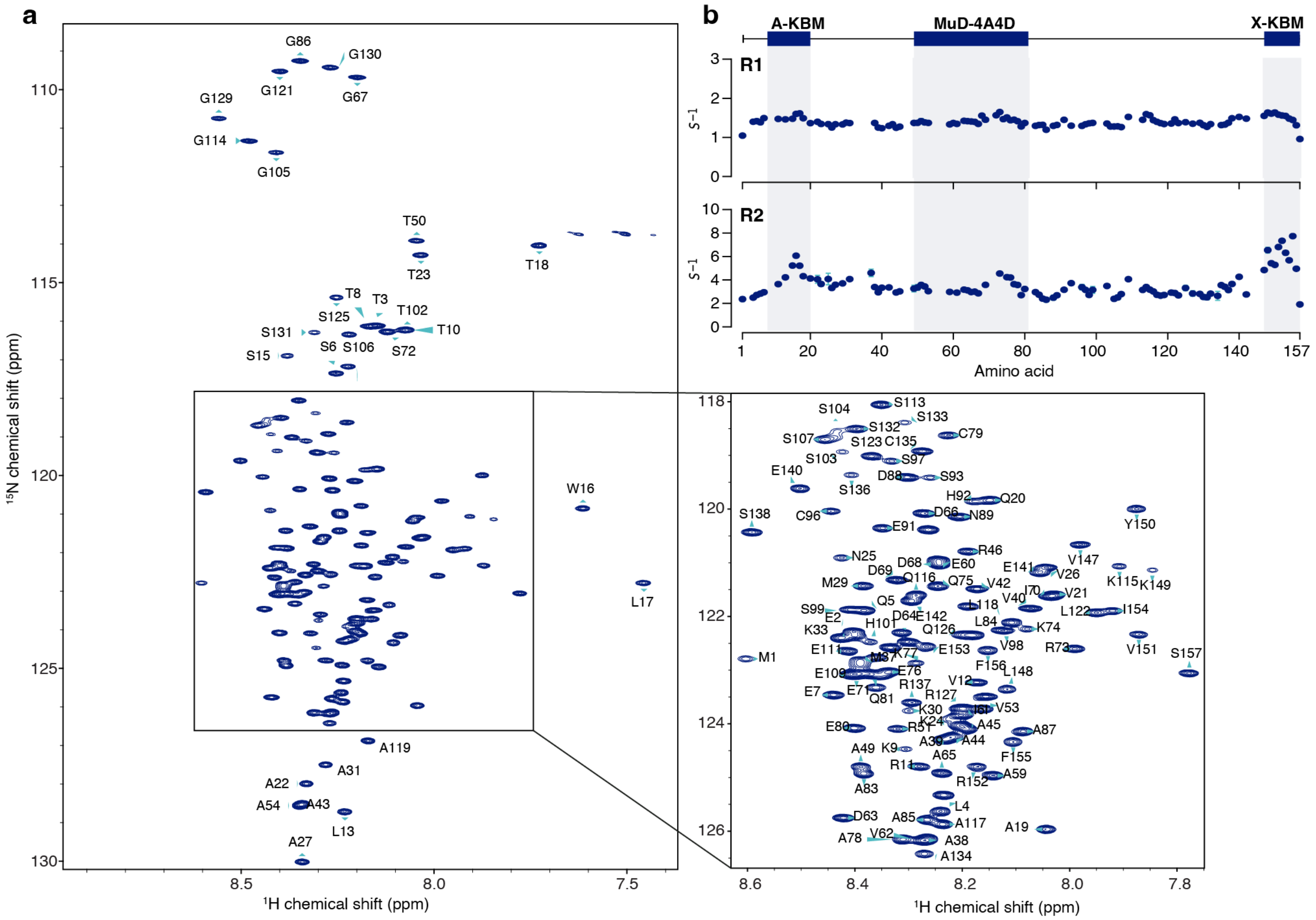


**Figure S7. MRI1 with 4A4D mutations is mostly disordered in the absence of urea.** (**a**) NMR resonance assignment of the $^{1}H$-$^{15}N$ HSQC spectrum of MRI1 4A4D with the enlarged inset. (**b**) Longitudinal relaxation rate ($R_1$) and transverse relaxation rate ($R_2$) of MRI1 4A4D. The experiments were measured on an 800 MHz spectrometer at 293 K. The error bars represent the uncertainty with one standard deviation, estimated either from the noise propagation, with n=1 independent experiment for each spectrum.

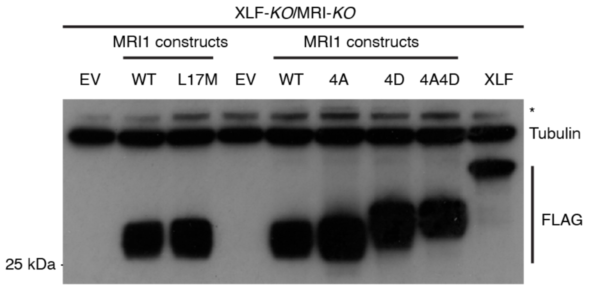


**Figure S8. Immunoblot of different MRI1 constructs and XLF in XLF and MRI1 double knockout cells.** All the proteins were probed with antibodies against Anti-Flag HRP. EV: empty vector, WT: wild-type, 4A: MRI1 $^{54}Y^{55}C^{56}M^{57}N$/AAAA, 4D: MRI1 $^{64}V^{66}L^{68}I^{69}L$/DDDD, 4A4D: MRI1: MRI1 $^{54}Y^{55}C^{56}M^{57}N$/AAAA-$^{64}V^{66}L^{68}I^{69}L$/DDDD.
